# Deep convolutional neural network with face identity recognition experience exhibits brain-like neural representations of personality traits

**DOI:** 10.1101/2024.03.28.587135

**Authors:** Wenlu Li, Jin Li, Tianzi Jiang

**Author notes:** Corresponding Authors: Tianzi Jiang.

## Abstract

Faces contain both identity and personality trait information. Previous studies have found that convolutional neural networks trained for face identity recognition spontaneously generate personality trait information. However, the successful classification of different personality traits does not necessarily mean that convolutional neural networks adopt brain-like representation mechanisms to achieve the same computational goals. Our study found that convolutional neural network with visual experience in face identity recognition (VGG-face) exhibited brain-like neural representations of personality traits, including coupling effects and confusion effects, while convolutional neural networks with the same network architecture but lacked visual experience for face identity recognition (VGG-16 and VGG-untrained) did not exhibit brain-like effects. In addition, compared to the VGG-16 and the VGG-untrained, the VGG-face exhibited higher similarity in neural representations with the human brain across all individual personality traits. In summary, these findings revealed the necessity of visual experience in face identity recognition for developing face personality traits judgment.

## INTRODUCTION

Faces are the most direct and prominent features to identify identities in our daily lives. Besides serving as primary cues for face identity recognition, faces also carry social information (*1*), for instance, we spontaneously infer the personality traits of other people based on their faces (*2, 3*). Therefore, faces simultaneously convey identity and personality trait information. However, it is currently unclear how face identity affects judgments of face personality traits.

An early influential distributed face processing model proposed that face identity information is primarily extracted by the ventral pathway of the face processing network, including the fusiform gyrus of the core system and the anterior temporal regions of the extended system, while personality trait information is mainly transmitted through the dorsal pathway of the face processing network to the medial temporal regions of the extended system (*4*). However, electrophysiological studies in monkeys and humans suggested that the medial temporal lobe (including the amygdala and hippocampus) may be involved in processing both face identity and personality trait information simultaneously. Research utilizing single-neuron recordings in humans has found that the amygdala is correlated with subjective ratings of face personality traits rather than objective face features (*5*). Recent studies using single-neuron recordings and simultaneous eye-tracking further demonstrated that certain neurons in the human amygdala and hippocampus encoded the spatial representation of face personality traits (*6, 7*). On the other hand, the amygdala and hippocampus also play crucial roles in face identity recognition tasks. Researchers recorded neural activity in the amygdala of monkeys viewing images of monkey faces, human faces, and objects, and found that the majority of neurons (64%) exhibited selective responses to face identity (*8*). Another study analyzed over 200 neurons in the amygdala of neurosurgical patients and found that during face-viewing tasks, 14% of amygdala neurons exhibited a significant main effect for face identity (*9*), consistent with previous findings from amygdala recordings in humans (*10, 11*). Recent research also discovered individual neurons in the monkey hippocampus showing selective responses when presented with faces of specific identities (*12*), providing information about face identity (*13*). An early study reported that a subset of neurons in the human medial temporal lobe was selectively activated by images of specific face identities (*14, 15*). Direct intracranial recordings in 50 human subjects found face-selective neural activity in the anterior of the medial temporal lobe and the entorhinal cortex, with the collaborative effort of these two regions facilitating face identity recognition (*16*). These studies suggest that the medial temporal lobe is involved in processing both face personality trait information and face identity information simultaneously. Therefore, we proposed a hypothesis that there is a mutual dependence between these two types of face information in the human brain. For example, the development of face personality trait judgment may depend on visual experiences of face identity recognition Due to ethical concerns, the need for long-term tracking, and the difficulty in controlling the visual experiences of participants (e.g., face identity recognition experience), studying the developmental mechanisms of the brain has become challenging in human developmental research. Bio-inspired artificial neural networks provide an effective model to address this issue. In recent years, an increasing number of studies have used artificial neural networks to simulate cognitive processing in the human brain. Specifically, research combining deep convolutional neural networks (DCNNs) with cognitive neuroscience has identified similar structural and functional characteristics between artificial and biological systems (*17*). Researchers have demonstrated a high degree of similarity between the hierarchical structure of convolutional neural networks and the ventral visual pathway in primates (*18, 19*). Moreover, the feature space in the deep layer of DCNNs serves as a useful model for simulating neural processes in the medial temporal lobe of the human brain. For example, researchers have demonstrated that features extracted from natural face images using DCNNs can be used to observe the features encoded by neuronal activity in the medial temporal lobe utilized in face identity recognition (*20*). On the other hand, generative computational models can be utilized to manipulate the process of face personality trait judgment in humans (*21, 22*). For instance, computer graphics models driven by human-rated scores can generate faces that elicit specific personality trait judgments. Other studies indicate that for a limited set of features, DCNNs can learn a direct mapping from face images to personality trait judgments without explicitly controlling or understanding the underlying two-dimensional or three-dimensional face structure (*23*). Furthermore, the outputs of DCNNs trained on different tasks can be used to predict human-made personality trait judgments. For example, a study utilized features from networks trained on object recognition, face recognition, and face landmark localization tasks to predict human-made personality trait judgments. The results revealed that features from networks trained on object recognition achieved the highest correlation between actual and predicted face personality trait judgments, demonstrating the widespread availability of visual feature information across different images (*24*). These studies suggest that DCNNs serve as effective computational models for investigating cognitive tasks related to face processing. Therefore, we hypothesize that if a DCNN simulating human face personality trait judgment requires visual experience in face identity recognition, it indicates that face personality trait information may depend on face identity information. Conversely, if it does not require such experience, it suggests that the processing of face identity information and face personality trait information may be separate.

Previous studies have found that DCNNs trained in face identity recognition contribute to the judgment of face personality traits (*24, 25*). However, when computational methods are used to explore cognitive functions in the human brain, successful classification of different personality traits only suggests a weak equivalence between DCNNs and humans in terms of input-output behavior and does not imply that DCNNs utilize brain-like representation to achieve the same computational goals (*26*). To validate the hypothesis that DCNNs share representation with the human brain, additional correlations should be established between computational models and the brain to verify stronger equivalence, such as their neural representational similarity (*27–29*). Therefore, in this study, we analyzed the similarity in the neural representations of face personality traits between DCNNs trained on face identity recognition in the medial temporal lobe of the human brain, to investigate whether brain-like judgment of face personality traits requires visual experience in face identity recognition.

One model used in this study was a typical DCNN pre-trained for face identity recognition (hereinafter referred to as VGG-face). If VGG-face can simulate the interdependence between face identity recognition and personality trait judgment in the human brain, it should spontaneously generate units that exhibit significant correlations between personality trait ratings, i.e., personality trait units. Furthermore, to test whether these personality trait units responded in a brain-like manner, it was necessary to compute their similarity to the neural representations of face personality traits in the human brain. Next, to validate that the brain-like neural representations of face personality traits formed by the neural network are not due to general object recognition experience or network architecture, two additional DCNNs were introduced: VGG-16, which has almost the same architecture as VGG-face but was trained for natural image classification, and untrained VGG-Face (hereinafter referred to as VGG-untrained), which has the same architecture as VGG-face but lacks any visual experience, with its weights randomly initialized. Comparing these three DCNNs will demonstrate whether brain-like neural representations of face personality traits require visual experience in face identity recognition, general object recognition, or merely depend on network architecture.

## RESULTS

### The emergence of personality trait units in the human brain and DCNNs

We first investigated the spontaneous emergence of personality trait units in both the human brain and DCNNs. Each participant viewed 550 face images and performed a simple one-back task (Fig. 1A). Each face image was displayed only once, with 50 randomly selected images repeated. Trials with repeated images were excluded from subsequent analyses. In the human brain, the firing rate of each unit was defined as the average firing rate within the time window of 0.25 to 1.25 seconds after baseline correction. A total of 1577 units with an average firing rate greater than or equal to 0.15 Hz were included in subsequent analysis, of which 753 were from the amygdala and 824 were from the hippocampus. Each DCNN (VGG-face, VGG-16 and VGG-untrained) comprised a feature extraction network and a classification network, with the last layer of the feature extraction network being the conv5-3 layer containing 100,352 units (Fig. 1B). 500 face images viewed by participants served as inputs to these DCNNs. Since the last layer of the feature extraction network (conv5-3) represents the highest-level representation among all convolutional layers and has the largest receptive field (*27, 30*), personality trait units were extracted from the conv5-3 layer. A unit was considered to be selective to personality traits if its outputs exhibited a significant correlation with any of the 8 personality trait ratings (p < 0.05, Spearman correlation) and showed no selectivity towards face identity (p ≥ 0.05, univariate ANOVA). The results revealed the spontaneous emergence of selective units for each personality trait in both the human brain and DCNNs, with some units exhibiting selectivity for more than one trait. Fig. 1C illustrates an example of a “warm” unit from the human brain, where the x-axis represents the z-scored firing rates and the y-axis represents the z-scored “warm” ratings, with gray dots representing individual images and the black line indicating the linear regression function between firing rates and ratings. The number of personality trait units in the human brain and the conv5-3 layer of DCNNs was presented in Table 1. In summary, the human brain contains 403 personality trait units (25.6%). Among the three DCNNs, the VGG-face exhibits the highest number of personality trait units, totaling 15,479 (15.4%), followed by VGG-16 with 4,333 personality trait units (4.3%), and the VGG-untrained with the lowest count of 2,303 personality trait units (2.3%).

**Fig. 1.**
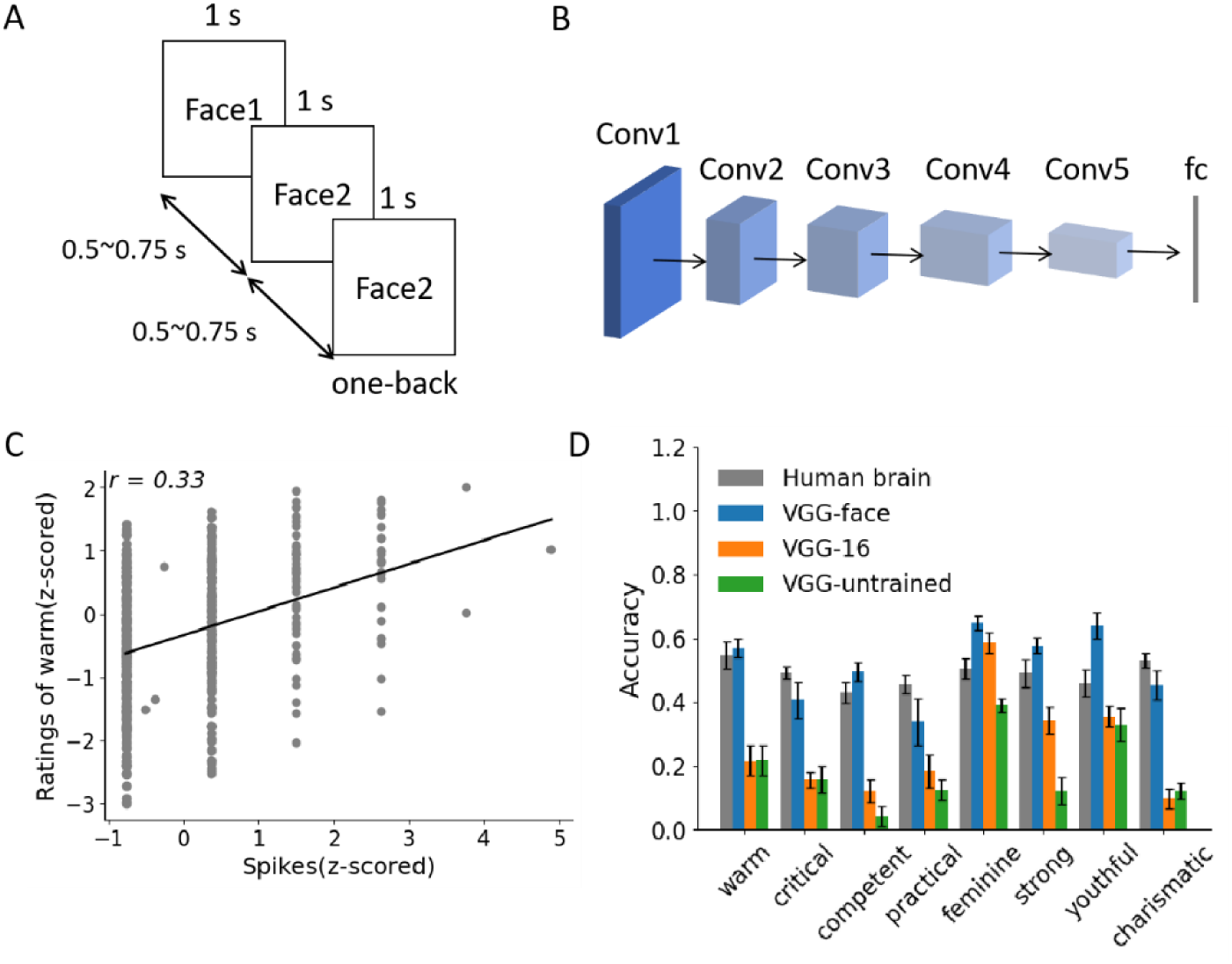
Experimental paradigm, DCNN architecture, and personality trait unit. **(A)** Experimental paradigm of human participants. Each participant viewed 550 face images and performed a simple one-back task. Each face image was presented for 1 second, with a stimulus interval (blank screen) of 0.5 to 0.75 seconds between images. Participants were required to press a key if the current face image matched the previous one. Trials with repeated images were excluded from subsequent analyses. **(B)** The architecture of DCNN. These convolution layers form a feature extraction network, converting images into target-oriented high-level representations, while the last fully connected layers form a classification network, categorizing images by converting high-level representations into classification probabilities. **(C)** An example of a “warm” unit in the human brain. This unit showed a significant positive correlation between firing rates (x-axis) and ratings of warm (y-axis). Gray dots represent individual images and the black line indicates the linear regression function between firing rates and ratings. **(D)** The prediction accuracy of ratings by the neural representations of personality traits in the human brain and DCNNs using linear regression with 10-fold cross-validation. The accuracy was measured by the Spearman correlation coefficient between the predicted ratings and the actual ratings. The error bars represent the standard error of accuracy across iterations.

**Table 1.** Number of personality trait units.

|  | Warm | Critical | Competent | Practical | Feminine | Strong | Youthful | Charismatic | Total |
| --- | --- | --- | --- | --- | --- | --- | --- | --- | --- |
| Human<br>brain | 68 | 73 | 65 | 83 | 87 | 81 | 90 | 77 | 403 |
| VGG-face | 6381 | 2633 | 2909 | 2916 | 6066 | 4130 | 3665 | 2885 | 15479 |
| VGG-16 | 1397 | 767 | 519 | 553 | 2468 | 933 | 1973 | 950 | 4333 |
| VGG-untrained | 561 | 506 | 572 | 542 | 1012 | 523 | 800 | 590 | 2303 |

The group activation of units for each personality trait forms a neural representation of that personality trait. For instance, there were 68 “warm” units in the human brain, and the outputs of these 68 units to all 500 face images constitute the “warm” neural representation (i.e., a matrix of shape 68×500 in math). In DCNNs, since the unit number of each personality trait exceeds 500, and there was considerable variation in the unit numbers across different personality traits, we performed Principal Component Analysis (PCA) on each personality trait neural representation to mitigate the impact of unit quantity on neural representations. To test the personality trait information provided by the neural representations of these units, we calculated their accuracy in predicting personality trait ratings using linear regression with 10-fold cross-validation. In each iteration, the accuracy was measured by the Spearman correlation coefficient between the predicted personality trait ratings and the actual ratings. Fig. 1D illustrates the prediction accuracy of ratings by the neural representations in the human brain and these DCNNs. The results showed that the human brain and VGG-face demonstrated higher prediction accuracy, indicating that the neural representations of personality traits of the human brain and the VGG-face provided more face personality trait information.

### DCNN with face identity recognition experience exhibited brain-like neural representations of personality traits

To investigate whether neural representations of personality traits in the human brain require domain-specific visual experience, such as face identity recognition or general object recognition experience, or solely rely on the architecture of DCNN, we computed the personality trait coupling matrices and confusion matrices for the human brain and each DCNN. Fig. 2 shows the schematic diagram of calculating the coupling effect and confusion effect (see Materials and Methods for details).

**Fig. 2.**
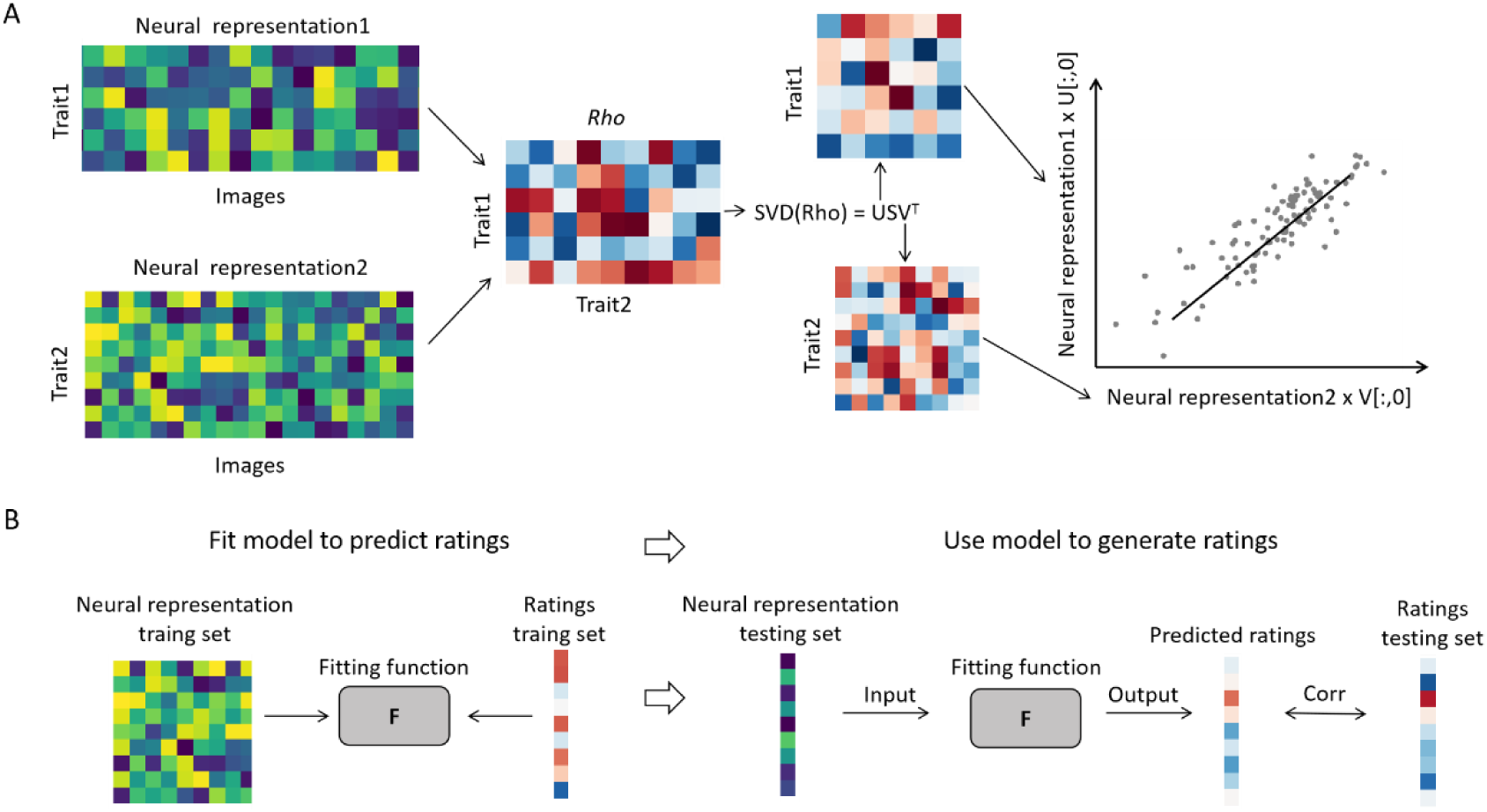
Overview of the procedure for calculating the coupling effect and confusion effect of personality trait. **(A)** Personality trait coupling effect. Correlations of pairs of personality trait neural representations were computed by decomposing the rank-correlation matrix of the two neural representations. Singular value decomposition (SVD) of the rank-correlation matrix results in neural weights per latent variable. The weights of the first latent variable (first column of each singular vector matrix) were used to reduce the dimensionality of neural representations (the number of personality trait units) into scores for each image. This method measures the latent estimate of across-image coupling between the two neural representations of personality traits. **(B)** Personality trait confusion effect. A linear regression function (F) was developed to predict ratings using neural representations of personality traits. The performance of the function was quantified using the Spearman correlation coefficient between the predicted ratings and the actual ratings was calculated with 10-fold cross-validation. This method measures the information provided by each personality trait neural representation for predicting different personality trait ratings.

The personality trait coupling matrices for the human brain and each DCNN are shown in Fig. 3A, B and C, where each element in the personality trait coupling matrix represents the maximum coupling effect between two personality traits. Since the personality trait coupling matrix was symmetric with diagonal elements equal to 1, the upper triangular elements of the personality trait coupling matrix were extracted and the correlations between the human brain and each DCNN were calculated separately. The results revealed a significant correlation between the personality trait coupling matrices of the human brain and the VGG-face (*r* = 0.49, *p* = 0.009, Spearman correlation). However, there was no significant correlation between the human brain and the VGG-16 or the VGG-untrained (VGG-16: *r* = 0.27, *p* = 0.16; VGG-untrained: *r* = 0.35, *p* = 0.09; Spearman correlation). The personality trait confusion matrices for the human brain and each DCNN are shown in Fig. 3D, E and F, where each element in the matrix represents the accuracy of predicting personality trait ratings using personality trait neural representations. Since the personality trait confusion matrices were asymmetric, the correlations between these matrices were calculated using all elements. The results demonstrated a significant correlation between the personality trait confusion matrices of the human brain and the VGG-face (*r* = 0.65, *p* < 0.0001, Spearman correlation). The important thing was, that there was no significant correlation between the human brain and the VGG-16 or the VGG-untrained (VGG-16: *r* = 0.11, *p* = 0.38; VGG-untrained: *r* = 0.04, *p* = 0.73; Spearman correlation), further supporting the necessity of face identity recognition experience for the formation of brain-like personality trait neural representations.

**Fig. 3.**
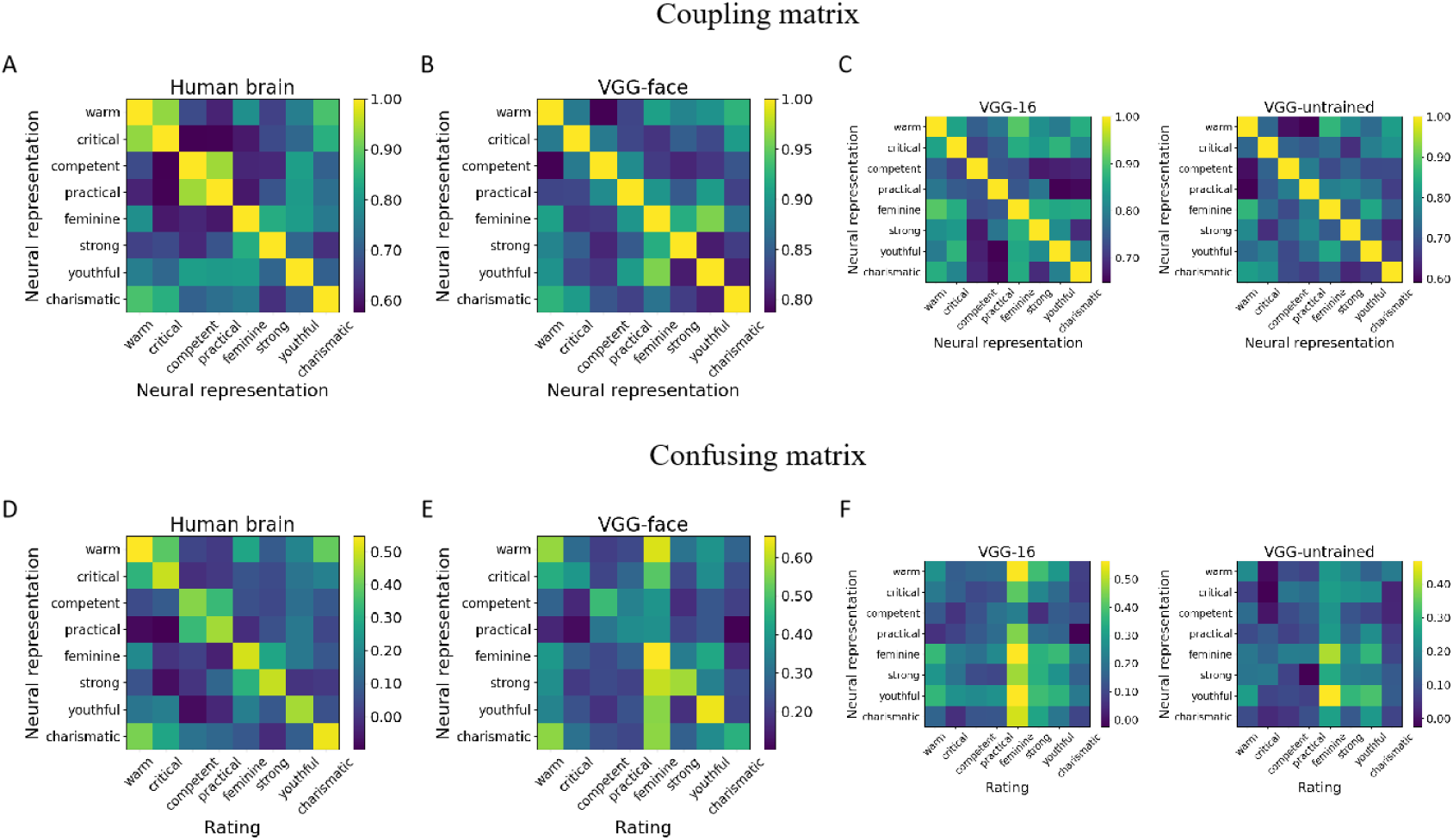
Personality trait matrices of the human brain and DCNNs. Coupling matrices of the (**A**) human brain, (**B**) VGG-face, (**C**) VGG-16 (left panel) and VGG-untrained (right panel). Confusion matrices of the (**D**) human brain, (**E**) VGG-face, (**F**) VGG-16 (left panel) and VGG-untrained (right panel).

### Brain-like neural representations of personality traits were not due to image pixel intensities or neural representations of face identity

To exclude the possibility that the similarity between the personality trait neural representations of the VGG-face and the human brain was due to the similarity in physical properties of the images, such as pixel intensities, we perturbed the pixels of each face image and calculated their personality trait neural representations of the VGG-face. This pixel perturbation method disrupted image features while preserving the pixel intensities. If the VGG-face exhibits brain-like effects on the perturbed pixel images, the similarity between the VGG-face and the human brain might be attributed to the pixel intensities of the images. Results indicated that there was no significant correlation between the personality trait coupling matrices and personality trait confusion matrices of the VGG-face and the human brain (personality trait coupling matrix: *r* = 0.27, *p* = 0.17; personality trait confusion matrix: *r* = 0.17, *p* = 0.19; Spearman correlation)) (Fig. 4A), suggesting that the brain-like personality trait neural representations exhibited by the VGG-face may be due to capturing structural features of face rather than image pixel intensities.

**Fig. 4.**
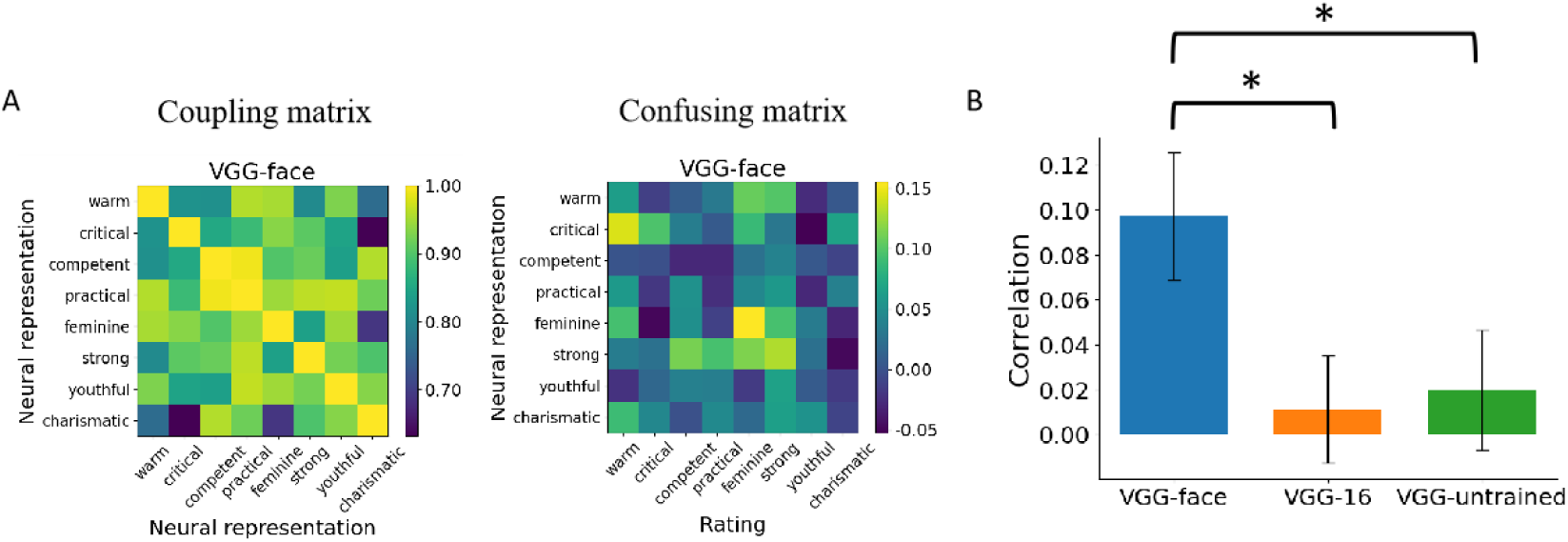
Brain-like neural representations of personality traits were not due to image pixel intensities or neural representations of face identity. (**A**) Personality trait coupling matrix (left panel) and personality trait confusion matrix (right panel) of the VGG-face for face images with perturbed pixels. (**B**) The correlations of the personality trait coupling matrices between DCNNs and the human brain for each face identity. Error bars represent the standard error of correlation coefficients across face identities. *p < 0.05.

Next, to rule out the possibility that the brain-like representations of the VGG-face were due to the neural representations of face identity, we calculated neural representation matrices of personality traits in the three DCNNs and the human brain for each identity. Since there were only 10 images for each identity, the accuracy of predicting personality trait ratings using neural representations for each identity was generally poor, so we only calculated the personality trait coupling matrix for each identity. For each DCNN and the human brain, there were 50 coupling matrices respectively, with each matrix only representing one face identity, so it does not contain information that can distinguish different identities. The correlations of the personality trait coupling matrices of the three DCNNs and the human brain were calculated separately. There were 50 correlation coefficients between each DCNN and the human brain, and each correlation coefficient represented the similarity between matrices with the same identity. The results showed that the similarities between VGG-face and the human brain were greater than those of the other two DCNNs (VGG-16: *p* = 0.03; VGG-untrained: *p* = 0.04; *N* = 50, paired *t-*test) (Fig. 4B), indicating the brain-like representations exhibited by VGG-face were not due to the neural representations of face identity.

### Local features and global configurations of faces provided different information for personality trait neural representations

Although both the personality trait coupling matrix and the personality trait confusion matrix measure the neural representations of personality traits, the information provided by these two matrices was not entirely identical. The personality trait coupling matrix reflected the degree of a monotonic correlation between neural representations, while the personality trait confusion matrix illustrated the ability of neural representations to predict personality trait ratings. To test whether different aspects of face features (global configurations or local features) provide different information, we applied two image processing methods to original face images: 1) to examine the information provided by the global configuration of faces (e.g., the distances between eyes, nose, and mouth relative to each other and the face contour) to neural representations, the images were divided into 5×5 patches and randomly recombined, thus disrupting the global configuration of the face but retaining the local features; 2) to examine the information provided by local face features (e.g., eyes, nose, and mouth) to neural representations, the images were blurred, thus disrupting the local face features but retaining the global configuration. The neural representations of the two types of processed images of the VGG-face were compared with the neural representations of the original face images of the human brain.

The results indicated that for the images disrupting the global configuration while retaining local face features, there was a significant correlation between the VGG-face and the human brain on the personality trait coupling matrices (*r* = 0.55, *p* = 0.002, Spearman correlation) (Fig. 5A). Conversely, there was no significant correlation between them on the personality trait confusion matrices (*r* = 0.08, *p* = 0.51, Spearman correlation) (Fig. 5B). This indicated that neural representations formed by global face configurations exhibit brain-like confusion correlations, contributing to the ability to predict personality trait ratings. On the other hand, for the images disrupting the local face features while retaining global configuration, a significant correlation was found between the VGG-face and the human brain on the personality trait confusion matrices (*r* = 0.52, *p* < 0.0001, Spearman correlation) (Fig. 5D). However, no significant correlation was observed between them on the personality trait coupling matrices (*r* = 0.07, *p* = 0.73, Spearman correlation) (Fig. 5C). This indicated that neural representations formed by local face features exhibit brain-like coupling correlations, contributing to the degree of monotonic correlations between neural representations.

**Fig. 5.**
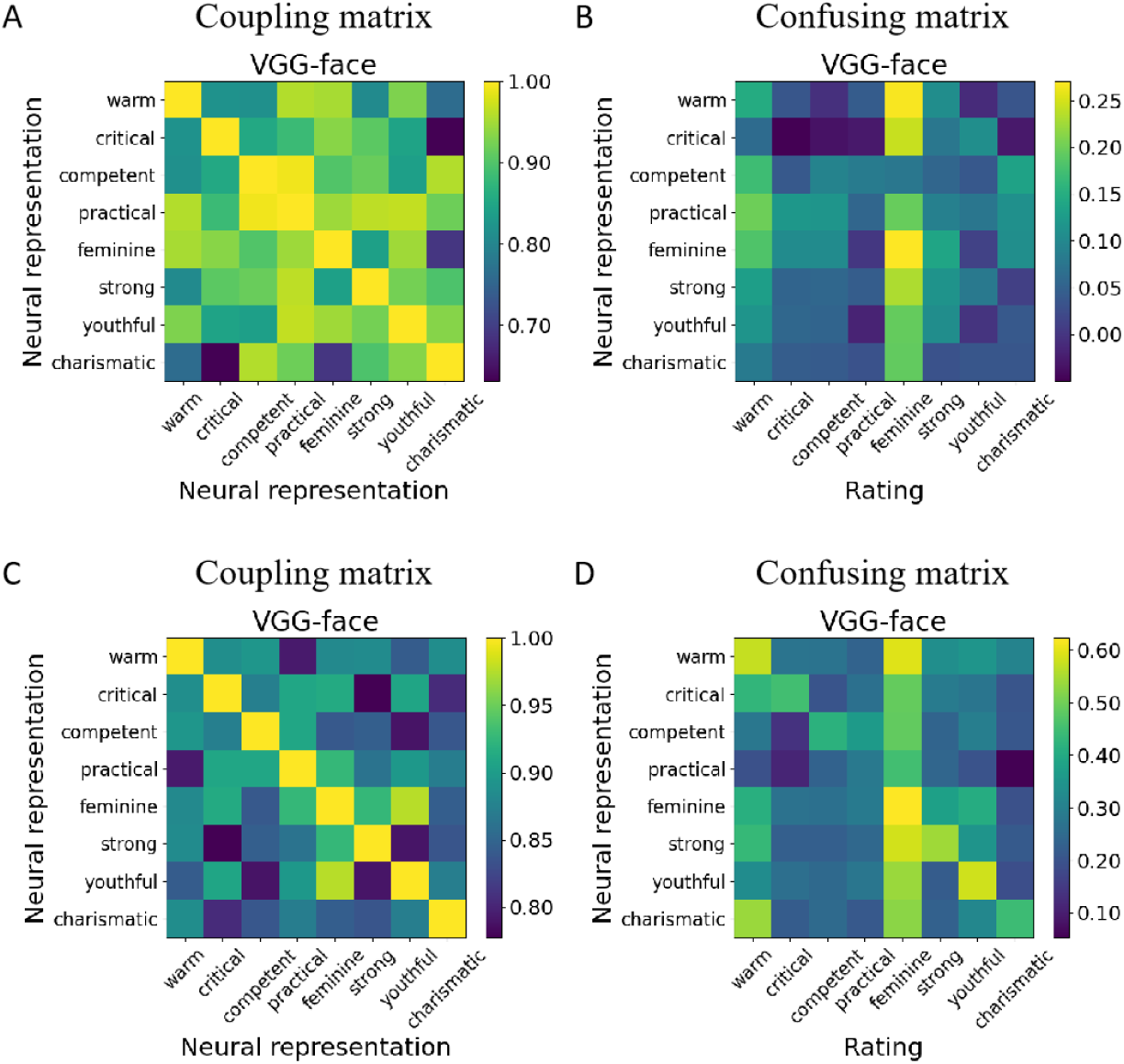
Personality trait matrices of the VGG-face for face images with disrupted global configuration or disrupted local features. (**A**) Personality trait coupling matrix and (**B**) personality trait confusion matrix of the VGG-face for face images with disrupted global configuration. (**C**) Personality trait coupling matrix and (**D**) personality trait confusion matrix of the VGG-face for face images with disrupted local features.

### VGG-face exhibited higher correlations with the human brain on individual personality trait representation

As both personality trait coupling matrices and personality trait confusion matrices measured correlations between personality traits, the similarities between these matrices reflected bidimensional correlations between these DCNNs and the human brain. To directly measure their correlations, we compared the correlations between these DCNNs and the human brain on individual personality traits. The calculation of correlation followed the algorithm of personality trait coupling matrices, with the difference being that while computing personality trait coupling matrices, variables represented neural representations of different personality traits within the individual DCNN or the human brain, whereas variables here represented the same personality trait neural representations between DCNNs and the human brain. Our hypothesis posited that if neural representations of personality traits require face identity recognition experience, then the VGG-face should exhibit higher correlations with the human brain compared to the VGG-16 and the VGG-untrained, particularly on individual personality traits. The results are shown in Fig. 6A, with dots representing neural representations of individual images, lines representing linear regression functions between DCNNs and the human brain. The Spearman correlation coefficients between the three DCNNs and the human brain on individual personality traits are shown in Fig. 6B. The VGG-face exhibited higher correlations with the human brain on all individual personality traits. Comparing the correlation coefficients between three DCNNs and the human brain on all personality traits, the results showed that the correlations between the VGG-face and the human brain were significantly higher than those of the VGG-16 and the VGG-untrained (VGG-16: *p* = 0.0045; VGG-untrained: *p* = 0.0003; *N* = 8, paired *t*-test), and the correlations between the VGG-16 and the human brain were significantly higher than those of the VGG-untrained (*p* = 0.03, *N* = 8, paired *t*-test) (Fig. 6C).

**Fig. 6.**
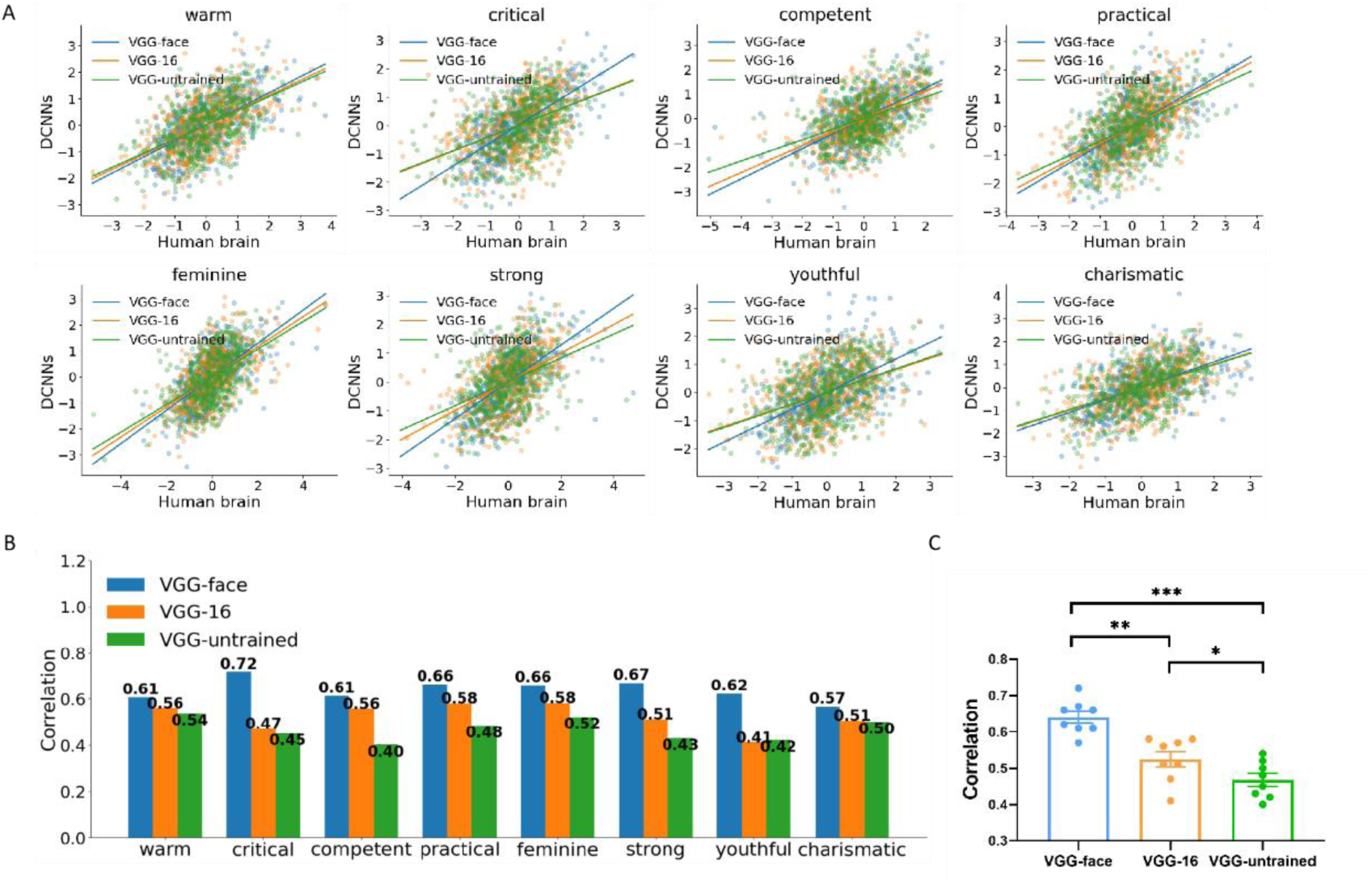
Correlations of neural representations between DCNNs and the human brain on individual personality traits. (**A**) Linear fitting of neural representations between DCNNs and the human brain on individual personality traits, with dots representing neural representations of individual images. (**B**) Bar graph of correlations between DCNNs and the human brain on individual personality trait neural representations. (**C**) Correlations between DCNNs and the human brain on all personality traits. *p < 0.05, **p < 0.01, ***p < 0.001.

## DISCUSSION

A core challenge in cognitive neuroscience is understanding how visual experiences shape neural representations in the brain. However, research into the development of human cognition faces various constraints, such as ethical concerns, recruitment difficulties, the need for long-term tracking, and the challenge of controlling the visual experiences of participants (e.g., face identity recognition experience). These limitations make it difficult to study the developmental mechanisms of the brain. Consequently, an increasing number of studies are turning to deep neural networks as powerful tools for investigating human cognitive development. In addition to exploring the weak equivalence between deep neural networks and humans in input-output behavior, researchers can establish similar neural representations between them through domain-specific training experiences.

This study identified units that selectively responded to personality traits in the medial temporal lobe (amygdala and hippocampus) of the human brain. Consistent with this finding, an earlier study found the amygdala of the human brain to be a necessary brain region for judging face personality traits, aiding in the retrieval of face personality trait-related knowledge (*31*). Recent research has revealed that the population of neurons in the medial temporal lobe encodes the personality trait space of faces, and this encoding is independent of face familiarity, identity or expression. Thus, the neural activity of the amygdala and hippocampus may be a universal neural mechanism encoding face personality traits. Furthermore, our study found that DCNN with face identity recognition visual experience (VGG-face), DCNN with only general object recognition visual experience (VGG-16), and DCNN with no visual experience (VGG-untrained) all spontaneously generated units selective to face personality traits. The phenomenon of face personality trait units spontaneously emerging in DCNNs without face visual experience is explicable. Previous research has found that DCNNs can gain sensitivity to abstract natural features by exposure to irrelevant natural visual stimuli (*32*), or even through randomly distributed weights without any visual experience (*33*). For example, units selective to face were found in randomly initialized networks, and these units exhibited some features observed in macaques, enabling untrained networks to perform face detection tasks. Similarly, another study also found face-selective units in untrained DCNNs with features similar to those of one-month-old macaques (*34*). These findings collectively suggested that random feedforward connections in untrained DCNNs may be sufficient to initialize primitive visual selectivity. Although all three DCNNs spontaneously generated units selective to face personality traits, the VGG-face had the most face personality trait units and achieved higher accuracy in predicting personality trait ratings. This result is consistent with a recent study indicating that DCNNs trained for face identity recognition facilitate the inference of personality traits from faces (*25*).

Further investigation reveals that only the personality trait neural representations in the VGG-face exhibit similarities with the human brain in terms of coupling effects, confusion effects, and higher similarities on all individual personality traits. The structures of the three DCNNs are identical, thus their differences in personality trait representations stem from different visual experiences. Therefore, the brain-like personality trait neural representations in the VGG-face may arise from face identity recognition visual experience. This may be because judging personality traits requires not only perceiving face features but also highly abstract social cognitive concepts, while using only generic natural features may struggle to extract abstract social information. These results emphasize the necessity of visual experience in face identity recognition for shaping judgments of face personality traits. Some studies have shown that using computer-generated models to create different face structures affects human personality trait judgments predictably (*22*). Since face structures also contribute to face identity recognition, we speculate that different face processing tasks may interact with each other through face structures.

Additionally, our findings indicate that global face configuration local face features provide different information for personality trait neural representations: global face configuration contributes to the ability to predict personality trait ratings based on neural representations, while local face features contribute to the degree of monotonic correlations between different personality trait neural representations. Previous research has shown that face identity recognition also involves the processing of both local face features and global configuration, which may play different roles in the face recognition process. For example, an fMRI study demonstrated that local face features and global configuration elicit responses in different brain regions (*35*), supporting the notion that local face features and global configuration provide different information for personality trait representations. Therefore, considering the mutual influence of face identity and personality traits, it is reasonable that the local face features and global configuration may play different roles in the judgment of face personality traits.

Early cognitive models of face processing suggested that the extraction of face identity information and personality trait information originates from different stages of face processing, processed through distinct pathways (*1*). The neural cognitive model proposed by Haxby et al. further elucidated the distinct neural anatomical circuits for these two types of information (*4*). Specifically, face identity information is primarily extracted by the ventral pathway of the face processing network, including the fusiform gyrus of the core system and the anterior temporal regions of the extended system, while personality trait information is mainly transmitted through the dorsal pathway of the face processing network to the medial temporal regions of the extended system. However, the results of this study indicate that face identity and face personality traits can form unified representations computationally. This suggests the possibility that there may also be a pathway between the ventral pathway and the medial temporal regions in the Haxby face processing model, where face identity information is transmitted from the ventral pathway to the medial temporal regions, supporting the formation of neural representations for face personality traits.

In summary, the results of this study demonstrate that VGG-face shares similar neural representations of face personality traits with the human brain, revealing the necessity of face identity recognition visual experience in the development of face personality trait judgments. Following exposure to face identity recognition tasks, the brain not only extracts identity features but also abstract social cognitive information, underscoring the contribution of acquired face identity recognition visual experience to the formation of face personality trait representations.

## MATERIALS AND METHODS

### Human brain data

The single-neuron recordings in the human brain for this study were obtained from a publicly available dataset (*36*). This dataset comprises recordings from 2082 neurons across 12 participants, with 996 neurons from the amygdala and 1086 neurons from the hippocampus. The participants were individuals undergoing intracranial electrode placement for the treatment of refractory epilepsy. All participants voluntarily participated and provided written informed consent in accordance with procedures approved by the Institutional Review Board of West Virginia University.

### Experimental paradigm

Each participant viewed 550 face images and performed a simple one-back task. The face images were obtained from the CelebA dataset, from which 50 identities (33 males) were selected, with each identity having 10 images, totaling 500 face images. These face images varied in expressions, angles, gaze directions, and background lighting, with some faces featuring accessories like sunglasses and hats. During the experiment, each face image was presented for 1 second, with a stimulus interval (blank screen) of 0.5 to 0.75 seconds between images. In the one-back task, participants were required to press a key if the current face image matched the previous one. Each face image was displayed only once, with 50 randomly selected images repeated, resulting in a total of 550 images. The purpose of the one-back task was to maintain the gaze of participants on the screen and allow participants to focus on face features, while avoiding potential biases towards specific face features (compared to requiring participants to judge specific face features). Trials with repeated images were excluded from subsequent analyses.

### Electrophysiology recording

Electrodes were implanted in the amygdala and hippocampus of patients with pharmacologically refractory epilepsy and single-neuron signals were recorded. Each electrode has eight microwires with a length of 40 microns, one of which was the reference signal. Signals were sampled at 32 kHz using bipolar broadband recording (0.1-9000 Hz) and analyzed offline using the Neuralynx system. The signal was then filtered using a 300-3000 Hz bandpass filter with 0 phase lag, and the discharges were sequenced using a semi-automatic template matching algorithm (*37*).

### Neuronal spikes

For each neuron, the average firing rate within the time window of 0.25 to 1.25 seconds after stimulus onset was considered as the response to the presented stimulus. Baseline correction was performed by subtracting the average firing rate within the time window of -0.25 to 0 seconds before stimulus onset. If the average firing rate after the stimulus presentation was lower than the average firing rate during baseline, the firing rate after the stimulus presentation was set to 0. Neurons with an average firing rate greater than or equal to 0.15 Hz were included in subsequent analysis.

### Rating of personality traits

Ratings from 439 typical individuals (283 males, with a mean age of 26.10 years) on eight personality traits for each face image: warm, critical, competent, practical, feminine, strong, youthful, and charismatic were collected from the Prolific online research platform. Ratings were measured on a 7-point scale ranging from 1 (not at all) to 7 (very apparent). The rating experiment was divided into 10 modules, with each module consisting of 50 face images, each from a different identity. Each individual provided a rating block for each personality trait. Three exclusion criteria were used for unfit ratings: 1) Trial exclusion: trials with response times less than 0.1 seconds or greater than 5 seconds; 2) Block exclusion: blocks with over 30% of trials excluded or with fewer than three different ratings; 3) Individual exclusion: individuals with more than three excluded blocks. Based on the above criteria, ratings for personality traits from 24 individuals and 7882 trials were excluded from further analysis.

### Deep convolutional neural networks

The pre-trained VGG-Face (www.robots.ox.ac.uk/~vgg/software/vgg_face/) has been trained on a dataset containing 2.6 million face images, enabling it to recognize 2622 identities (*38*). This network consists of 13 convolution layers and 3 fully connected layers. The first 13 convolution layers are distributed across 5 blocks. In the first 2 blocks, each block contains 2 consecutive convolution layers followed by a max-pooling layer. In the subsequent 3 blocks, each contains 3 consecutive convolution layers followed by a max-pooling layer. The first 13 convolution layers form a feature extraction network, converting images into target-oriented high-level representations, while the last 3 fully connected layers form a classification network, categorizing images by converting high-level representations into classification probabilities (*39*). The architecture of VGG-16 is identical to the VGG-face, except for the last fully connected layer, which contains 1000 units for classifying 1000 object categories. The VGG-untrained retains the same architecture as the VGG-face, but lacks training experience and was assigned random connection weights (initialized through Xavier normal distribution).

### Personality trait coupling matrix

The personality trait coupling matrix was obtained by computing the coupling effect between pairs of personality trait neural representations. The coupling effect was quantified using latent modeling (*40*). Specifically, this involves calculating the rank correlation matrix (*Rho*) between two personality trait neural representations, followed by Singular Value Decomposition (SVD) of the rank correlation matrix. This decomposition yields the left singular vector matrix *U*, the right singular vector matrix *V^T^*, and the diagonal matrix of singular values *S*:

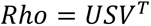

The singular vector matrix contains orthogonal latent variables. By applying the first column of each singular vector matrix to the neural representations and calculating the correlation between the transformed neural representations, the maximum correlation between the two neural representations of personality traits was measured.

### Personality trait confusion matrix

The personality trait confusion matrix was obtained by using the neural representation to predict its own and other personality trait ratings. Prediction accuracy was quantified using 10-fold cross-validation. Specifically, the dataset was first randomly divided into 10 equal parts (each containing 50 images), where 9 parts were used as training sets, and the remaining part was used as a test set. A linear regression function was then trained using the personality trait neural representations and ratings in the training set, and the personality trait neural representations in the test set were inputted into this function to obtain predicted ratings. Finally, the Spearman correlation coefficient between the predicted ratings and the actual ratings was calculated to measure the performance of the fitting model. This process was repeated 10 times to ensure predictions were obtained for all samples. The correlation coefficients obtained from all iterations were averaged to obtain the performance of the model on the entire dataset. This method measures the information provided by each personality trait neural representation for predicting different personality trait ratings.

## Notes

### Competing Interest Statement

The authors have declared no competing interest.

